# Thermal Tolerance is linked with Virulence in a Fish Pathogen

**DOI:** 10.1101/188185

**Authors:** Roghaieh Ashrafi, Matthieu Bruneaux, Lotta-Riina Sundberg, Katja Pulkkinen, Janne Valkonen, Tarmo Ketola

## Abstract

Although increase in temperatures may boost the number of pathogens, a complex process involving the interaction of a susceptible host, a virulent strain, and environmental factors would influence disease virulence in unpredictable ways. Here we explored if the virulence of an environmentally growing opportunistic fish pathogen, *Flavobacterium columnare*, would be malleable to evolutionary changes via correlated selection on thermal tolerance. Virulence among the strains increased over years, but tolerance to higher temperatures was associated with reduced virulence. Our results suggest that observed increase in frequency of columnaris epidemics over the last decade is most likely associated with increased length of growing season, or other time dependent change in environment, rather than increased regional average temperatures. Our results also indicate that most virulent bacteria had weaker ability to tolerate outside host environments, which suggest trade-off between more obligate pathogen behaviour and ability to grow outside host.

## INTRODUCTION

Climate projections suggest that changing climate not only affects the average temperature but also the occurrence of extreme and variable temperatures ^1^. Such changes in climate alter extinction risks, provoke range shifts and cause selection pressure to favour genotypes that are adapted to cope with these new environments ^2-4^. Microbes, many of which have the capacity to be or become pathogens, are expected to adapt rapidly. Global warming may benefit many bacterial species, since they will face milder winter months resulting in greater overwintering success, increased numbers of generations and, thus, higher pathogen densities to damage hosts ^5,6^. Environmentally growing opportunistic pathogens, in contrast to obligate (fully host-dependent) pathogens, can utilize outside host resources, making them very sensitive to selection pressures outside the host ^7^. Therefore, predicting the effect of climate warming on environmental opportunistic bacteria with life cycles both outside and inside the host present a particular challenge because pathogen fitness in both environments may be differentially affected by temperature ^8^. Although the ability to stay alive in the environment, e.g. as inactive spores, has been linked with high virulence ^9,10^, pathogens can also evolve towards a more benign virulence since investments in resource acquisition and defence in the outside-environments can trade off with traits connected to virulence ^11,12^. Previous studies suggest that higher temperatures select genotypes that tolerate hotter temperatures, whereas fluctuations in temperature should select for more generalist genotypes with improved tolerance to extreme temperatures ^13-18^. Nevertheless, it has remained unclear how climate warming might affect growth parameters in environmentally growing opportunistic pathogens, and how this correlates with their potential to cause disease.

Understanding the selection pressures underlying the evolution of virulence in outside-host environments is crucial in the current context of climate change, especially for diseases affecting world food production. *Flavobacterium columnare*, the etiological agent of columnaris disease in farmed fish, is an opportunistic fish pathogen which severely impacts freshwater aquaculture worldwide ^19,20^. Specifically, this bacterium can cause infections both in cold and warm water fish species ^21^. The temperature range in which it can grow actively is approximately 15 to 35°C ^19^. Previous work on this bacterium in the context of global warming and even a number of virulent pathogens has focused mainly on long-term empirical data examining the relationship between mean ambient temperature and disease prevalence ^22,23^. Analysis of more than 20 years’ worth of data has showed a significant positive effect of mean water temperature on the prevalence of columnaris disease at two fish farms ^22^. However, it is still unclear if climate change will impact the thermal performance of this bacterium at long term by selecting more thermo-tolerant strains and if such changes may have any effect on bacterial virulence. This is important information for regions where climate change is expected to be most severe, such as Finland where average annual temperature is predicted to rise nearly twice as fast as the average temperature for the whole globe 24.

Thermal tolerance is usually depicted via thermal performance curves (TPC) composed from the measured performance of a genotype in different thermal environments. Assuming that thermal performance curves obtained from measurements done in constant environments can be used to predict how genotypes survive under fluctuations ^25-27^, adaptation to fluctuating environments could occur either via overall elevated TPC or via broadened TPC ^16,17,28^. The key ecophysiological parameters that characterize thermal performance curves are the critical thermal thresholds which represent the lower (CT_min_) and upper (CT_max_) temperatures at which performance (e.g. growth or yield of bacteria) is zero, the optimum temperature (T_opt_) at which performance is maximal, and the maximum value of performance itself (µ_max_). In addition to these parameters, variation in TPC can also be characterized using principal component analysis in order to identify the main patterns of performance variation among the genotypes ^29,30^. However, the latter method has been rarely applied to thermal performance data in bacteria.

In the present study, we measured bacterial growth at five different temperatures (spanning from 17 to 32 that matches typical summer growth season in Finland, and in the near future) in order to characterize the temperature dependence of maximum biomass (hereafter yield) in 49 *F. columnare* isolates collected across Finland. Based on this data, we examined (i) variation of thermal performance in isolates, and (ii) the link between thermal performance and bacterial virulence, using virulence data measured in a separate experiment. We showed that Finnish isolates differed in maximum yield and limits of thermal range, and that their good tolerance to high temperatures was linked to lowered virulence.

## MATERIAL AND METHODS

### F. columnare strains and culture conditions

We used 49 Finnish *F. columnare* isolates for which genotypes were previously determined by the conventional MLST method using six loci ^31^ (Supplementary Table 1). All strains, belonging to broadly defined genetic group characterized by good low-temperature tolerance (genomovar I) ^31,32^, were originally isolated from eight fish farms, from both Southern (approx. 65° N) and Northern (62°N) parts of Finland (Supplementary Table 1), from fish or tank water using standard culture methods with modified Shieh medium ^33^, Shieh medium supplemented with tobramycin ^34^, or AO-agar ^35^.

**Table 1.** Effect of MLST genotype, year of collection and location on strain thermal performance estimates. Marginal means are reported for each level of the qualitative variables (Genotype, Location) and slope is reported for the continuous variable Year. For variables which were removed from the final model, the reported values (in italics) are the ones obtained in the last step before they were removed during model selection.

|  | Estimate | Std error | F-value | (df1, df2) | P-value |
| --- | --- | --- | --- | --- | --- |
| **$\mu_{max}$** |  |  |  |  |  |
| **Genotype** |  |  | **18.7779** | **(3, 44)** | **<0.001** |
| C | 1.091 | 0.012 |  |  |  |
| E | 0.993 | 0.015 |  |  |  |
| G | 1.044 | 0.028 |  |  |  |
| A&H | 0.922 | 0.025 |  |  |  |
| *Location* |  |  | *1.8084* | *(1, 43)* | *0.186* |
| Northern | 1.035 | 0.020 |  |  |  |
| Southern | 1.004 | 0.012 |  |  |  |
| **Year** | -0.007 | 0.004 | 3.6614 | (1, 44) | 0.062 |
| **Topt** |  |  |  |  |  |
| **Genotype** |  |  | **3.2075** | **(3, 42)** | **0.033** |
| C | 26.118 | 0.164 |  |  |  |
| E | 25.762 | 0.184 |  |  |  |
| G | 24.922 | 0.367 |  |  |  |
| A&H | 25.660 | 0.300 |  |  |  |
| *Location* |  |  | *0.6578* | *(1, 41)* | *0.422* |
| Northern | 25.427 | 0.269 |  |  |  |
| Southern | 25.681 | 0.157 |  |  |  |
| *Year* | -0.028 | 0.050 | *0.3254* | *(1, 40)* | *0.572* |
| **CTlow** |  |  |  |  |  |
| **Genotype** |  |  | **5.2715** | **(3, 37)** | **0.004** |
| C | 17.577 | 0.178 |  |  |  |
| E | 16.511 | 0.212 |  |  |  |
| G | 17.028 | 0.397 |  |  |  |
| A&H | 16.668 | 0.459 |  |  |  |
| *Location* |  |  | *0.2220* | *(1, 35)* | *0.640* |
| Northern | 16.864 | 0.320 |  |  |  |
| Southern | 17.036 | 0.199 |  |  |  |
| *Year* | 0.057 | 0.055 | *1.0523* | *(1, 36)* | *0.312* |
| **CThigh** |  |  |  |  |  |
| **Genotype** |  |  | **7.3380** | **(3, 45)** | **<0.001** |
| C | 31.570 | 0.167 |  |  |  |
| E | 32.471 | 0.200 |  |  |  |
| G | 31.107 | 0.400 |  |  |  |
| A&H | 32.722 | 0.326 |  |  |  |
| *Location* |  |  | *0.5217* | *(1, 44)* | *0.474* |
| Northern | 32.144 | 0.285 |  |  |  |
| Southern | 31.903 | 0.171 |  |  |  |
| *Year* | 0.005 | 0.054 | *0.0093* | *(1, 43)* | *0.924* |
| **TPB** |  |  |  |  |  |
| **Genotype** |  |  | **7.8427** | **(3, 37)** | **<0.001** |
| C | 13.994 | 0.275 |  |  |  |
| E | 15.970 | 0.328 |  |  |  |
| G | 14.079 | 0.614 |  |  |  |
| A&H | 15.437 | 0.709 |  |  |  |
| *Location* |  |  | *0.4048* | *(1, 35)* | *0.529* |
| Northern | 15.080 | 0.496 |  |  |  |
| Southern | 14.721 | 0.308 |  |  |  |
| *Year* | -0.069 | 0.086 | *0.6543* | *(1, 36)* | *0.424* |

### Thermal performance measurements

Bacterial isolates were grown overnight in modified Shieh medium under constant agitation (120 rpm) in room temperature and further sub-cultured to fresh medium in ratio of 1:10 for another 16-18 h under the same conditions. Sterile 15 ml tubes containing 5.5 ml of bacterial culture were centrifuged for 5 min in 4°C at 3500 *g*, after which the supernatant was discarded. 240 µl of concentrated bacterial culture was mixed with 60 µl of 10% of glycerol and 10% of fetal calf serum mixture on 100-well Bioscreen C^®^ plate in a randomized order and stored at −80°C. Prior to growth measurements, bacterial isolates were inoculated to a new Bioscreen plate containing 400 µl fresh modified Shieh medium in each well directly from the frozen Bioscreen plate using heat-sterilized cryo-replicator (Enzyscreen B.V., Haarlem, Netherlands ^36^). After 24 h incubation at 25°C, inoculums of 40 µL of individual bacterial strains from these pre-cultures were distributed into a Bioscreen plate containing 400 µL of fresh modified Shieh medium in each well for the growth measurements. Growth experiments were run simultaneously in duplicate in two 100-well plates in a Bioscreen C spectrophotometer (Oy Growth Curves Ab Ltd, Finland) over two to eight days depending on the experimental temperature, at five different temperatures (17°, 20°, 22°, 29° and 32°C). The bacteria were cultured without shaking, and optical density (OD) measurements were performed at 5-minute intervals (absorbance at 600 nm). The growth curves were analysed as described in Ketola *et al.* ^16^ to estimate maximum growth rate and maximum biomass or yield. Maximum yield estimates were more robust than maximum growth estimates due to the sensitivity of the latter to noise in the early phase of the growth curves. Consequently, we chose to use maximum yield as a measure for strain performance at a given temperature.

Two alternative approaches were used to analyse the thermal performance data: (i) curve fitting followed by estimation of TPC parameters (CT_min_, CT_max_, T_opt_, µ_max_) for each strain and (ii) principal component analysis (PCA) on the discrete performance measurements.

### Thermal performance curve fitting and parameter estimation

We used the TableCurve 2D software (version 5.01; Systat Software, Inc. 2002) to select a set of 1960 candidate equations to describe the relationship between yield and temperature. Using data from a subset of experimental strains, all equations using two-and three-term functions, with intercept available in the TableCurve 2D library, were fitted and the resulting fits with large R^2^ values were visually inspected. Candidate equations were selected based on the fulfilment of the following criteria to ensure a biologically meaningful fit: (i) “bell-shaped” curve with maximum yield occurring within the experimental thermal range, (ii) mostly concave curve (i.e. curves with several and clear local maximums in the experimental thermal range were discarded, but slight bumps were allowed), (iii) extrapolation outside the experimental thermal range predicted decreased performance (i.e the behaviour of the curve outside the experimental thermal range was consistent with biological expectations). In the end, the following 6 equations were chosen as candidates for a plausible model of the relationship between temperature (x) and performance (y):

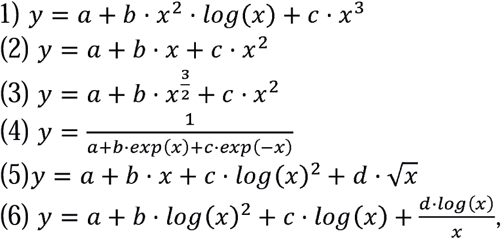

where a, b, c and d are strain-specific curve parameters. The average R^2^ values across the 49 strains used in this study were greater than 0.98 for each of those equations.

For each strain, a weighted-average thermal performance curve was built after fitting those six candidate equations, where AIC values were used to calculate a strain-specific weight for each of the six equations according to the formula:

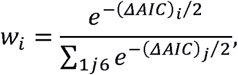

where *w_i_*is the weight assigned to the i^th^ equation and *(ΔAIC)_i_is* the difference between the AIC of the i^th^ equation and the lowest AIC among the six equations for this strain. While acknowledging that our procedure for the selection of candidate equations might introduce some subjectivity in the choice of candidate curves, keeping 6 different candidate equations and producing a weighted-average model based on their AIC values allowed for a variety of shapes in the final fitted curves with an overall good quality of fit, as shown in Supplementary Figure 1.

The obtained average thermal performance curves were used to determine maximum performance µ_max_ and optimal temperature T_opt_. We decided not to extrapolate unreasonably the thermal performance curves to determine CT_min_ values, but instead chose to estimate thermal ranges by calculating for each strain the temperatures at which its TPC reached half its maximum performance, hereafter CT_50_/_low_ and CT_50_/_high_. Growth at lower temperatures falls gradually and the growth in this species is already un-measurable at 15°C, causing estimation inaccuracy if curve fitting and in estimating CT min. Thermal performance breadth (TPB) was defined as the difference between the estimated CT_50/high_ and CT_50/low_. A visual inspection of the fitted curves was performed to remove CT_50/low_ (for 8 strains) and T_opt_ values (for 3 strains) which were unreliable given the shape of the fit for a particular strain (e.g. very flat plateau at µ_max_, unreliable extrapolation for CT_50/low_), resulting in 41 strains with all TPC parameters.

### PCA on yield measurements

Since PCA is sensitive to outlier data points departing from normal distributions, we visually inspected normal quantile-quantile plots of the yield data to identify and remove three outliers out of 49 strains prior to PCA. PCA was performed using the covariance matrix of maximum yields in five temperatures. Since the outliers were removed only to avoid unduly affecting the PCA by their departure from normality but were otherwise biologically meaningful, the coordinates of all 49 strains along each principal component (PC) were then calculated based on the PCA loadings obtained from the subset of 46 strains. In order to facilitate the biological interpretation of the patterns of variation described by each PC, we predicted the TPC of hypothetical strains located at the extreme boundaries of the 95% range of the coordinates of experimental strains along each PC using the inverse of the PCA matrix.

### Virulence assay

A virulence experiment was conducted according to the Finnish Act on the Use of Animals for Experimental Purposes, under permission ESAVI-2010-05569/Ym-23 granted for L-RS by the National Animal Experiment Board at the Regional State Administrative Agency for Southern Finland. Virulence of the 49 bacterial strains was assessed in an experiment using zebra fish (*Danio rerio*). The fish were infected using bacterial cultures grown overnight in fresh modified Shieh medium and adjusted at 4x10^5^ colony-forming units (CFU) mL^−1^. Ten fish per bacterial strain were individually challenged in 500 ml of water by adding 500 µl of adjusted bacterial culture directly into the experimental aquaria. The water temperature was maintained at 25°C during all experiments, which is close to the mean T_opt_ of the strains used. It has been shown than the zebra fish can be used as a reliable model in virulence experiments since it shares the temperature optimum of this pathogen ^37^. Aquaria containing fish were randomly placed on shelves in the experimental room to reduce the differences between aquaria. This infection method has been shown to produce a rapid onset of disease in fish, bringing out strain differences ^37,38^. As a control, 10 fish were individually exposed to 500 µl of sterile Shieh medium. Disease signs and fish morbidity were monitored in two hour intervals for 97 hours. Morbid fish that had lost their natural swimming buoyancy, and which did not respond to external stimuli, were considered dead and removed from the experiment, and euthanatized by cutting the spinal cord to avoid the suffering of the fish. The experiment was conducted according to the Finnish Act on the Use of Animals for Experimental Purposes, under permission granted by the National Animal Experiment Board at the Regional State Administrative Agency for Southern Finland for L-RS.

### Statistical analyses of thermal performance data

The effects of MSLT genotype group (categorical variable, 4 levels), year of strain isolation (continuous variable) and geographical location (categorical variable, 2 levels: Northern and Southern Finland) on thermal performance were assessed using model selection starting from a full linear model specified as:

> Performance = intercept + β_g_.Group + β_y_.Year + β_l_.Location + residuals

where Performance was either one of the thermal performance curve parameters estimated from curve fitting (µ_max_, T_opt_, CT_50/low_, CT_50/high_ or TPB) or coordinates along one of the principal components of interest (PC1, PC2 or PC3). No interaction between Group and Year or Group and Location was included in the starting model due to the imbalanced distribution of strains from different MSLT genotype groups across the years or across the geographical range of our study. Model selection was performed iteratively: at each step, variables were dropped one at a time and the significance of the change in fit for each dropped variable was tested using a Chi-square test (function *drop1* in R). If the highest p-value for significance of change in fit was greater than 0.10, the corresponding variable was dropped from the model and the next selection step was performed; otherwise model selection was stopped.

### Statistical analyses of virulence data

Since the vast majority of death events occurred early in the virulence assay, virulence data was analysed by considering fish survival as a binary variable (death/survival). The effects of explanatory variables on fish death were estimated using generalized linear mixed models (binomial family) with a logit link function and using strain identity as a random factor. Two full models differing in how they incorporated thermal performance as an explanatory variable (using either (i) PCs or (ii) TPC parameters) were used as starting models. The fixed effects used in those two initial models were:

> (i) MLST genotype, year, location, PC1, PC2, and PC3 (49 strains)

> (ii) MLST genotype, year, location, µ_max_, T_opt_, and CT_50/high_ (46 strains)

CT_50/low_ and TPB were not included in full model due to colinearity with CT high (Figure 1.).

**Figure 1.**
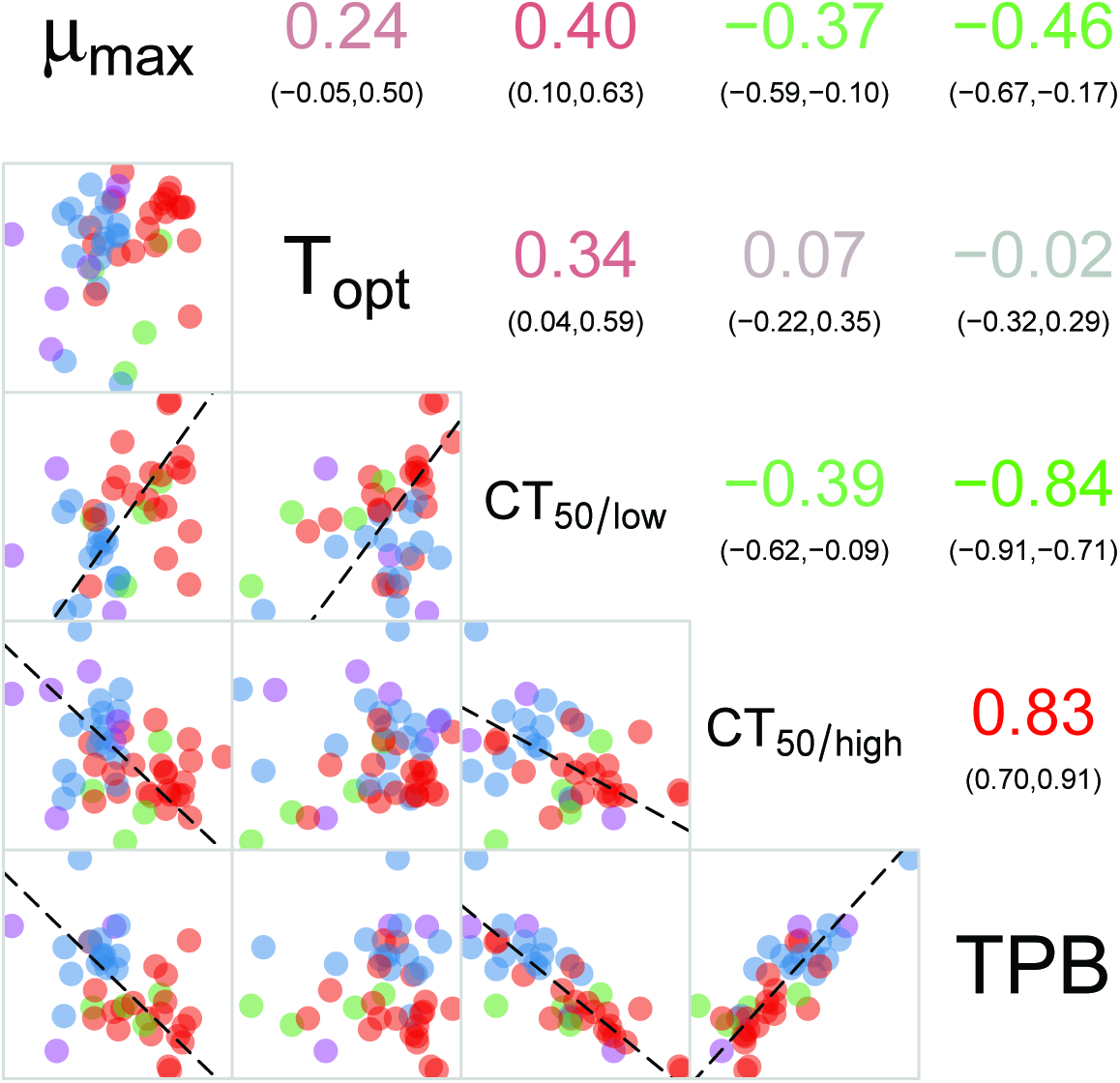
Correlogram for thermal performance parameter estimates. Upper triangle, Pearson∙s product moment correlation coefficients between pairs of variables. The numbers in brackets indicate the 95% confidence interval. Lower triangle, scatter plot between pairs of variables. When the Pearson∙s product moment correlation coefficient is significant (p-value < 0.05), a dashed line indicates the ranked major axis. For the upper triangle, the color coding on a green to red scale matches the correlation coefficient value (−1, green; 0, grey; +1; red). For the lower triangle, colors match MLST genotype: red, purple, green, and blue are for genotypes C, E, G and A&H, respectively.

Models were fitted using the *glmer* function from the *lme4* package in R. Model selection was performed starting from each of the full models and testing the effect of removing one variable out of MLST genotype, year and location at a time, and testing for the significance of the change in fit with a likelihood-ratio test (function *drop1* in R). At each step, the variable with the highest p-value for the significance of change in fit was dropped if this p-value was greater than 0.10. We did not remove any thermal performance variable included in the initial starting models since we wanted to consider all the thermal performance characteristics simultaneously in the models. We used the DHARMa package in R ^39^ to assess the correctness of the residuals.

## RESULTS

### Correlations between thermal performance curve parameters

TPC parameters were estimated from the AIC-weighted average curves for each of the 49 strains; due to uncertainty in estimated values for some fits, T_opt_ values were calculated for 46 strains, and CT_50/low_ values for 41 strains (Supplementary Figure 1, Supplementary Table 1). A correlogram was built to explore pairwise correlations between TPC parameters (Figure 1): CT_50/low_ and CT_50/high_ were negatively correlated, suggesting a gradient between narrow and wide thermal performance range (specialist-generalist gradient). T_opt_ was positively correlated with CT_50/low_ but not with CT_50/high_, which reflects a horizontal shift of the left-hand part of the TPC while maximum thermal tolerance would be more constrained. Finally, µ_max_ was positively correlated with CT_50/low_ and negatively correlated with CT_50/high_ and TPB; this might reflect a trade-off between increased tolerance to a larger range of temperatures and higher maximum performance.

### Principal components describing variation in thermal performance

We selected the first three principal components, which accounted for 93% of the variability of the yield measurements at 17, 22, 24, 29 and 32°C (Figure 2, Supplementary Table 2). Based on the predicted TPC along each PC (Figure 2), PC1 (describing 46% of the variation) is related to an increase in the thermal performance breadth by increasing yield at both extreme temperatures (17°C and 32°C) while the maximum performance is unchanged along this component. PC1 is thus mostly a generalist-specialist axis. PC2 (describing 30% of the variation) is characterized by a negative correlation between performance in cold and warm temperatures: PC2 can be seen as a cold adaptation/warm adaptation axis. PC3 (describing 17% of the variation) corresponds to a change in the maximum performance, negatively correlated with performance in the coldest temperature but unrelated to performance at the warmest temperature.

**Figure 2.**
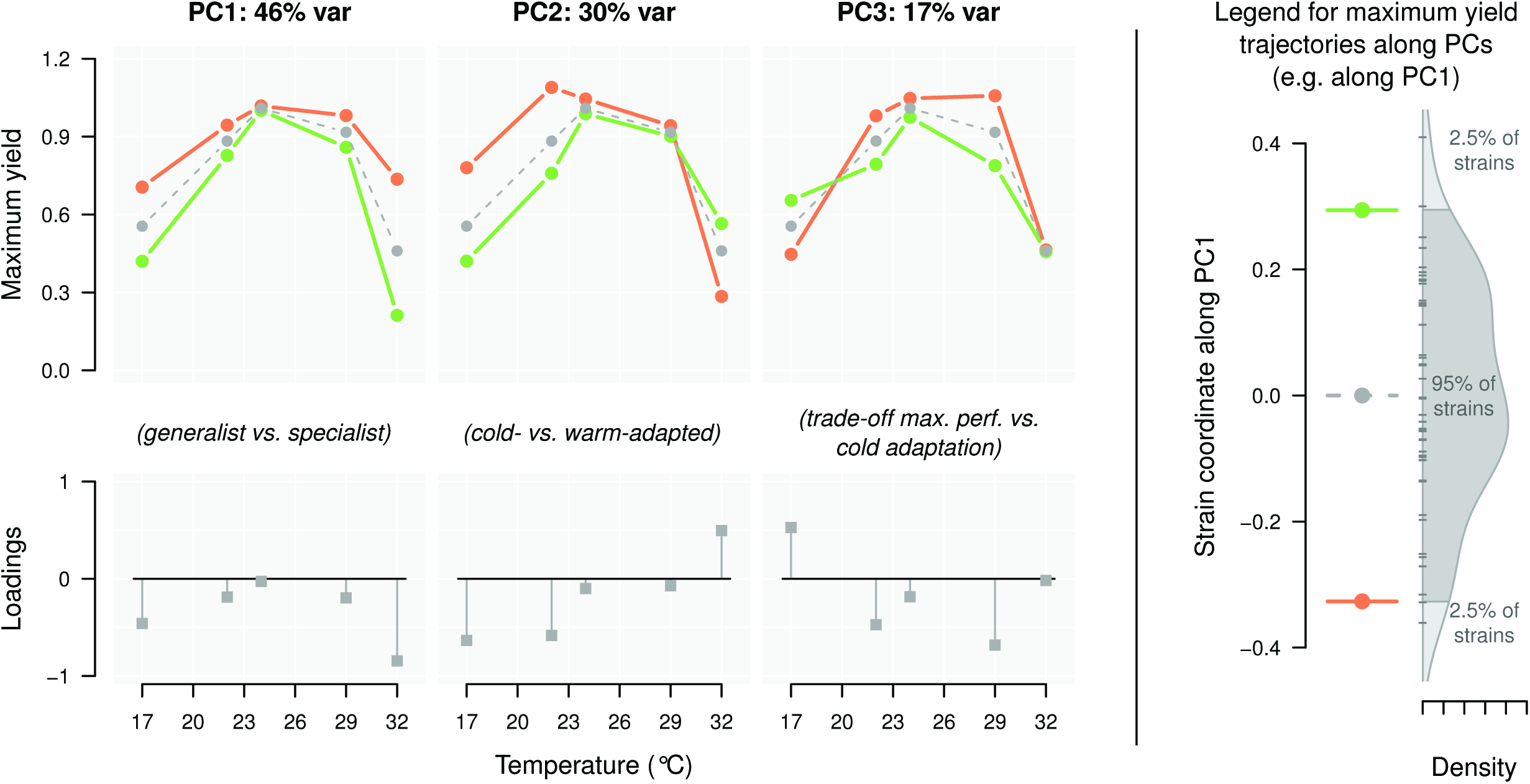
PCA results and their interpretation in terms of TPC patterns. Upper panels, prediction of TPC variation patterns along each of the first three PCs. Grey dashed line, average performance curve of the 49 strains used in this study. Orange and green lines, performance curves of hypothetical strains with coordinates at the lower and upper 95% quantiles, respectively, along each PC, as depicted in the explicative panel on the right. Lower panels, loadings for each original temperature on the first three PCs.

**Table 2.** Effect of MLST genotype, year of collection and location on strain coordinates along PCs. Marginal means are reported for each level of the qualitative variables (Genotype, Location) and slope is reported for the continuous variable Year. For variables, which were removed from the final model, the reported values (in italics) are the ones obtained in the last step before they were removed during model selection.

|  | Estimate | Std error | F-value | (df1, df2) | P-value |
| --- | --- | --- | --- | --- | --- |
| **PC1** |  |  |  |  |  |
| *Genotype* |  |  | 1.3983 | (3, 45) | 0.256 |
| *C* | -0.006 | 0.040 |  |  |  |
| *E* | 0.063 | 0.048 |  |  |  |
| *G* | -0.123 | 0.097 |  |  |  |
| *A&H* | 0.091 | 0.079 |  |  |  |
| *Location* |  |  | 2.1451 | (1, 44) | 0.150 |
| *Northern* | 0.092 | 0.068 |  |  |  |
| *Southern* | -0.025 | 0.041 |  |  |  |
| *Year* |  |  | 0.1342 | (1, 43) | 0.716 |
|  | -0.005 | 0.013 |  |  |  |
| **PC2** |  |  |  |  |  |
| *Genotype* |  |  | 0.8297 | (3, 43) | 0.485 |
| *C* | -0.030 | 0.040 |  |  |  |
| *E* | -0.037 | 0.059 |  |  |  |
| *G* | -0.146 | 0.093 |  |  |  |
| *A&H* | 0.038 | 0.079 |  |  |  |
| **Location** |  |  | 6.1481 | (1, 47) | 0.017 |
| Northern | -0.094 | 0.039 |  |  |  |
| Southern | 0.030 | 0.031 |  |  |  |
| *Year* |  |  | 0.8896 | (1, 46) | 0.351 |
|  | 0.009 | 0.010 |  |  |  |
| **PC3** |  |  |  |  |  |
| **Genotype** |  |  | 9.8268 | (3, 45) | <0.001 |
| C | -0.060 | 0.018 |  |  |  |
| E | 0.051 | 0.021 |  |  |  |
| G | 0.002 | 0.042 |  |  |  |
| A&H | 0.120 | 0.034 |  |  |  |
| *Location* |  |  | 1.6877 | (1, 44) | 0.201 |
| *Northern* | -0.005 | 0.030 |  |  |  |
| *Southern* | 0.040 | 0.018 |  |  |  |
| *Year* |  |  | 1.2299 | (1, 43) | 0.274 |
|  | 0.006 | 0.006 |  |  |  |

### Determinants of thermal performance

The MLST genotype affected all calculated thermal performance parameters (µ_max_, T_opt_, CT_50/low_, CT_50/high_ and TPB) (Table 1). Year effect was close to significance for µ_max_, with maximum performance decreasing slightly over the years (Table 1). Geographical location had no significant effect on any TPC parameter.

Location had a significant effect on PC2 coordinates, with Northern strains exhibiting lower PC2 values (Table 2), corresponding to cold adaptation (Figure 2). MLST genotype had a significant effect on PC3 coordinates (negative correlation between maximum performance and cold tolerance, Figure 2).

### Determinants of virulence

When the effect of thermal performance on virulence was analysed using TPC parameters estimated from curve fitting, 46 strains out of 49 could be included in the analysis due to missing values in T_opt_. Year of isolation had a positive effect on virulence (Figure 3B, Table 3).

**Figure 3.**
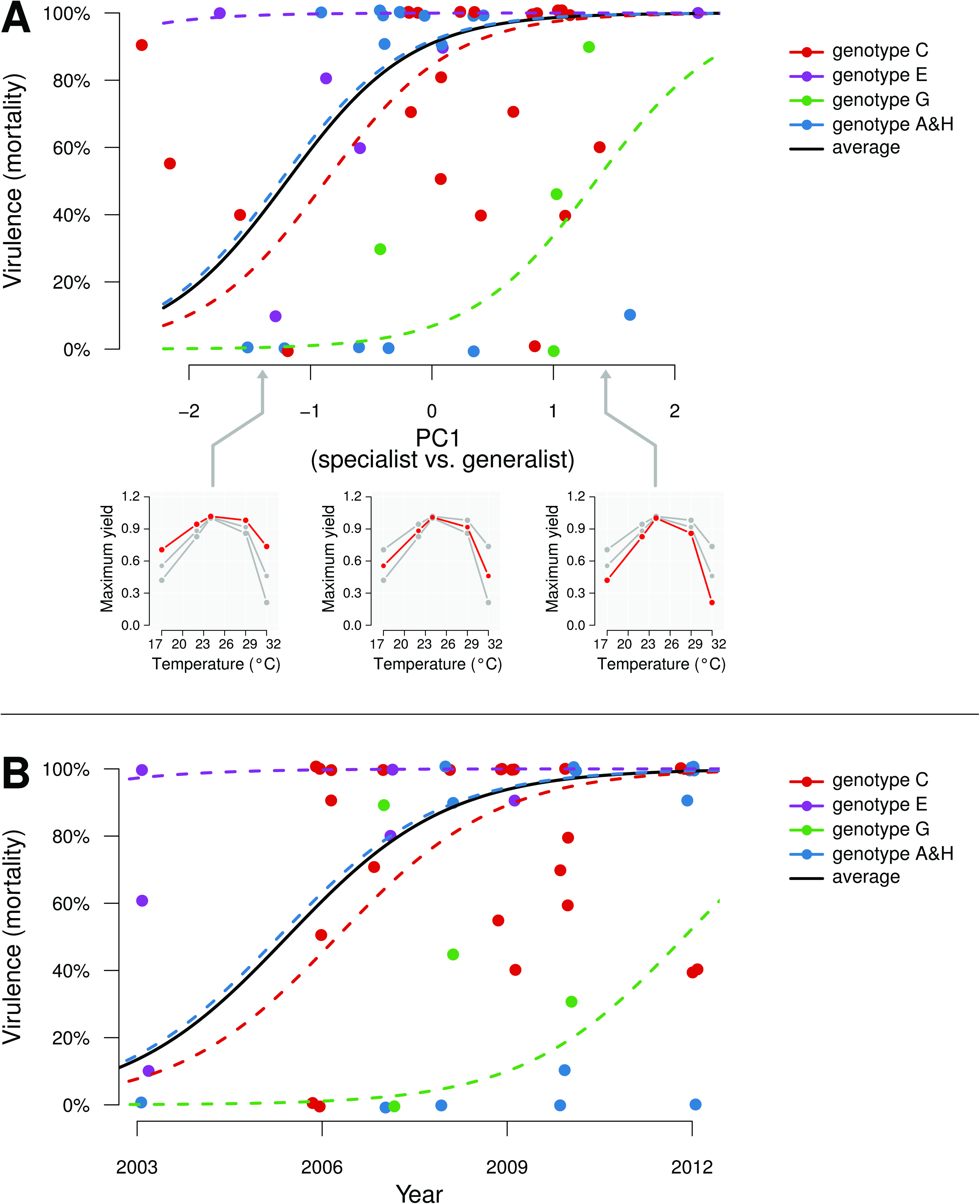
Effect of PC1 coordinate and year of collection on strain virulence. Each marker represents the average mortality observed for a given strain. Fitted curves are plotted using the GLMM results presented in Table 4. Colours represent genotype groups. The black fitted line correspond weight-averaged fixed effect estimates based on the abundance of genotype groups in our dataset. Panel A, effect of PC1 coordinate on strain virulence. The three sub-panels illustrate how TPC varies from specialist to generalist phenotype along PC1. Panel B, effect of year of collection on strain virulence.

**Table 3.** Effect of strain characteristics on virulence (using PCs coordinates). Model used in R: death ~ genotype + year + location + PC1 + PC2 + PC3 + (1|strain), with a binomial family distribution and a logit link function. Marginal means and standard errors are reported for each different level of qualitative variables (Genotype and Location) in the original response scale (pDeath) while slope estimates and standard errors in the logit scale are reported for the continuous variables. For variables which were removed from the final model, the reported values (in italics) are the ones obtained in the last step before they were removed during model selection.

|  | Estimate | Std<br>error | Chi-square | Df | P-value |
| --- | --- | --- | --- | --- | --- |
| **Genotype** |  |  | 4.4056 | 3 | 0.221 |
| C | Pdeath=0.665 | 0.327 |  |  |  |
| E | Pdeath=0.907 | 0.135 |  |  |  |
| G | Pdeath=0.203 | 0.424 |  |  |  |
| A&H | Pdeath=1.000 | 0.001 |  |  |  |
| *Location* |  |  | *0.0303* | *1* | *0.862* |
| *Northern* | *Pdeath=0.945* | *0.098* |  |  |  |
| *Southern* | *Pdeath=0.923* | *0.083* |  |  |  |
| **Year** |  |  | 4.3022 | 1 | **0.038** |
|  | 2.366 | 1.141 |  |  |  |
| **µmax** |  |  | 0.4359 | 1 | 0.509 |
|  | 0.765 | 1.159 |  |  |  |
| **Topt** |  |  | 3.6112 | 1 | 0.057 |
|  | 1.765 | 0.929 |  |  |  |
| **CThigh** |  |  | **4.4010** | **1** | **0.036** |
|  | -2.588 | 1.234 |  |  |  |

Among analysed TPC parameters, only CT_50/high_ had an effect on virulence (Table 4): strains with higher tolerance to high temperatures were less virulent. When the effect of thermal performance on virulence was analysed using PCs (49 strains used), both year and PC1 coordinate had a significant effect on virulence: strains collected more recently were more virulent (similarly as observed using TPCs) and more generalist strains had lower virulence (Table 4, Figure 3A).

**Table 4.**
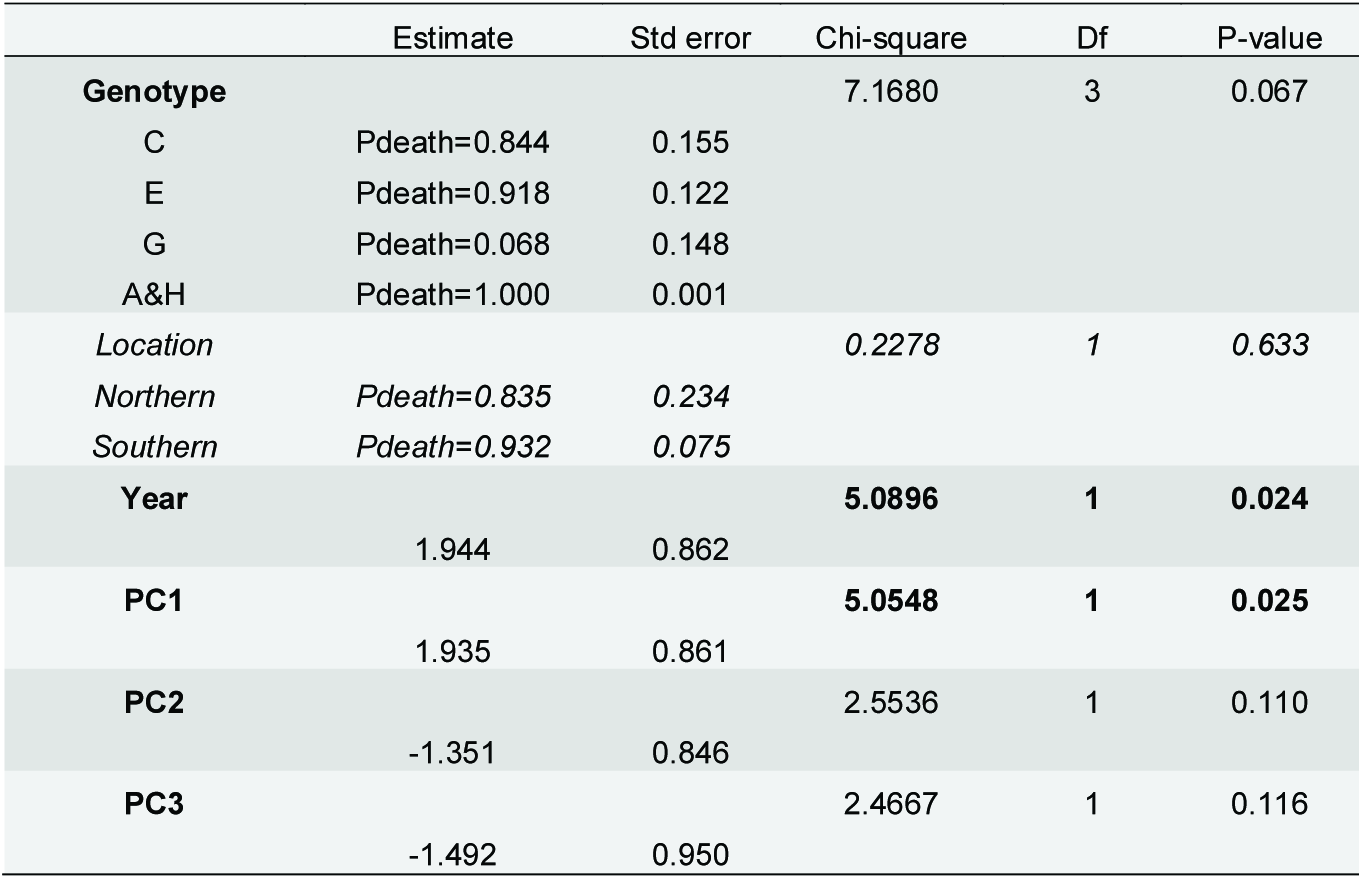
Effect of strain characteristics on virulence (using TPC estimates). Model used in R: death ~ genotype + year + location + µmax+ Topt+ Topt2+ CT50/low+ CT50/high + (1|strain), with a binomial family distribution and a logit link function. Continuous variables (Year, µmax, Topt, Topt2, CT50/low, CT50/high) were z-normalised. Marginal means and standard errors are reported for each different level of qualitative variables (Genotype and Location) in the original response scale (pDeath) while slope estimates and standard errors in the logit scale are reported for the continuous variables. For variables which were removed from the final model, the reported values (in italics) are the ones obtained in the last step before they were removed during model selection.

## DISCUSSION

There is a growing body of evidence indicating that some pathogens become more prevalent ^40,41^ and more virulent at warmer temperatures ^42^. For example, increased expression of virulence factors is correlated with increased temperature in *Vibrio* species ^43,44^. Although global warming may increase the number of virulent pathogens, pathogens with free-living stages and ectothermic hosts are particularly susceptible to changes in temperature because temperature can have complex and opposing effects on different parts of the pathogen life cycles ^8^. We explored if strains of an aquaculture-associated pathogen vary in their thermal performance, and if thermal performance was correlated with strain virulence. This type of information is crucial in predicting how future climate change scenarios could alter environmental pathogens’ virulence via correlated selection on their thermal performance. In theory, due to co-evolutionary shifts in both host and the pathogen, virulence is expected to decrease over time, as fitness of both populations is optimized. However, virulence is context-dependent, as both biotic factors such as host condition ^46^ and host density ^47^ and abiotic factors such as temperature ^48^ can influence virulence. For example, high environmental temperature could select for increased bacterial pathogenicity, in *Serratia marcescens* ^49^.

We characterized the temperature dependency of maximum biomass production (i.e. yield) of 49 isolates of *F. columnare* that were collected from eight different areas located across Finland between 2003 and 2012. We performed temperature performance curves (TPC) and used principal component analysis on raw performance measurements to assess the variation of thermal performance between strains. Our results revealed that despite northern location Finnish *F. columnare* typically have a rather high optimum temperature between 23.7 °C and 27.9°C and an upper critical temperature for yield between 30.1°C and 34.7 °C. Finnish lakes form predominantly closed and shallow basins (average depth about 7 metres) and surface waters may reach high temperatures in summer. Tolerance to high temperature might be necessary for inhabiting natural waters during summer since this bacterium has an environmental origin ^50^. Consistent with the idea that cold tolerance is a key element for survival and growth in high latitudes, isolates from Northern Finland were more tolerant to colder temperatures than isolates from Southern Finland (see: PC2 in Figure 2 and effect of location on PC2 in Table 2). Our findings are in agreement with previous studies showing that selection may favour higher performance in higher altitude/latitude environments to guarantee successful reproduction and transmission during short growth seasons ^45^.

On the other hand, ample amount of genotype dependent variation in all TPC parameters (Table 1, Figure 1) suggests thermal performance is strongly constrained by genetic background of the bacteria. These findings clearly indicate that thermal conditions can in principle have a strong effect on the genetic diversity of *F. columnare* in the environment, and therefore presumably also on disease dynamics.

One would expect that *F. columnare* with high optimum temperatures may lead to more epidemics in the future owing to climate change. Interestingly, virulence was negatively correlated with upper thermal tolerance (CT_50/high_) and with thermal generalism (Tables 3 and 4). This suggests that not higher temperatures or increased fluctuations associated with climate change should not select for higher virulence in this species. However, since the CT_50/high_ values are already very high (beyond 30°C) it is unlikely that any increase in temperatures in the near future would select for changes in CT_50/high_ in the strains isolated in Finland, due to clearly lower maximum temperatures in fish farms. Furthermore, as the epidemics become more frequent at the high temperatures, recurring antibiotic treatments during onset of epidemics leave little possibility for co-selection for virulence and performance in high temperatures to take place, especially when temperatures in the farming environment rarely exceed +25^°^C (supplementary Figure 3).

Guijarro *et al.* ^48^ showed that numbers of bacterial diseases in aquaculture, particularly those of freshwater, can occur at temperatures below bacterial optimal growth (optimum growth temperature for the fastest growth under laboratory conditions). Therefore, the constraints of maximum environmental temperature on virulence should be relatively limited. This view is supported by considering outbreaks of epidemics, which start to occur at farms when water temperatures exceed ca. 18°C, after which epidemics are often treated with antibiotics. Temperature records from a fish farm in Central Finland over the past few decades show that maximum temperatures during the outbreak season have not significantly increased, but rather the overall growth season length has increased (Figure S3). Still, our results show that Finnish *F. columnare* strains have become intrinsically more virulent in recent years, as evidenced in our experiments under controlled laboratory conditions where confounding effects such as increased environmental temperature, variable nutrient availability or variable host density were removed (year effect in Table 3 and Table 4 and figure 3-B) (see also^51^). Thus, yearly increases in virulence could be a consequence of increased growing season, due to intensified fish farming ^51^, or due to some other time associated change in environment.

We showed that maximum performance was overall negatively correlated with thermal performance breadth, suggesting a trade-off between generalism and high-performance specialism (Figure 1). However, we did not find that generalist genotypes with broader performance breadth would have lower biomass at optimum temperatures and hence opposing the classic generalist-specialist trade off hypothesis. It is noteworthy that theories are highly idealized and a “Jack of all temperatures” does not always have to be a master of none ^52^: genotypes can have broader thermal performance range without paying a visible performance cost at optimum conditions, but possibly involving a trade-off with other traits ^14,16,54^, such as virulence ^16,53^.

For environmentally growing opportunist pathogens, adaptations for more efficient exploitation of one growth environment could be expected to cause repercussions in pathogens ability for growth in the other environment ^7^, such as host. Alternatively, the presence of virulence factors in the bacteria is unnecessary during the planktonic state but essential for the infection process, helping bacteria to save energy by not expressing virulence genes until they sense they have entered the host environment ^48^. This could explain why more generalist strains with broader thermal performance breadth, were less virulent than more specialist ones (see: PC1 effect in Table 4 and Figure 3-A). Similarly, expression of virulence factors was found to lower the outside growth rate in *Salmonella typhimurium* ^53^ and adaptation to tolerate thermal fluctuations and predators have caused lowered virulence in experimental evolution settings^11,16,49,55^.

In conclusion, it seems that current problems with steadily increased severity of outbreaks and evolved virulence cannot be directly linked to increased mean temperature in fish farms and associated bacterial evolution. Still, the found clear genotype and location effects on several thermal tolerance parameters suggest that temperatures can play strong role in dictating which genotypes and clones of this important fish pathogen are successful in different thermal environments.

## ACKNOWLEDGMENT

We would like to thank Dr. Ilkka Kronholm and Dr. Elina Laanto for providing constructive comments and help in improving the contents of this paper. We would like to thank Dr. Päivi Rintamäki, Dr. Heidi Kunttu, MSc. Reetta Penttinen and Dr. Elina Laanto for donating bacterial isolates for this study. We would also like to thank MSc Jenni Marjakangas for valuable help in the lab. The authors want to thank Yrjö Lankinen from Savon Taimen for providing the temperature data from Tyyrinvirta. This work was supported by KONE foundation (Roghaieh Ashrafi via project “Constraints of evolutionary adaptation to climate change” to Tarmo Ketola), OLVI foundation (Roghaieh Ashrafi #201620393), the Jane and Aatos Erkko Foundation (Lotta-Riina Sundberg), Finnish Cultural Foundation (Katja Pulkkinen) and Academy of Finland (Lotta-Riina Sundberg #266879, Tarmo Ketola # 278751, Jouni Taskinen # 260704 to Katja Pulkkinen) and Centre of Excellence in Biological Interactions (#252411, Prof. Johanna Mappes) to Roghaieh Ashrafi, Lotta-Riina Sundberg, and Tarmo Ketola.

## SUPPLEMENTARY MATERIAL

**Sup. Figure S1. TPC fits**. Each plot represents maximum yield (y-axis) versus temperature (x-axis). The strain name is indicated in each panel. For each strain, the six candidate equations from TableCurve 2D were fitted (grey lines) and the resulting curves were averaged using AIC-weights (red lines). Thermal performance parameters (maximum yield, optimum temperature, CT_50/low_ and CT_50/high_) were determined from the average curve for each strain. Asterisks denote strains for which estimated values for either CT_50/low_ alone (*) or CT_50/low_ and T_opt_ (**) were deemed too unreliable and were set as missing values in downstream analyses.

**Sup. Figure S2. Evolution of seasonal variation in water temperature in a Finnish fish farm (Tyyrinvirta fish farm)**. The graphs show the evolution of (A) monthly average temperatures, (B) averages of three highest temperatures per month and (C) averages of three lowest temperatures per month. Red lines are fitted using a linear model within each month. The p-value for the significance of the year effect on the monthly values is reported for each month.

**Sup. Table S1. Strain information**. Site, year of isolation, source of isolation (fish or water), location of isolation, host species, sequence type (ST), genotype, maximum biomass (yield) estimates (µ_max_), optimal temperature (T_opt_), lower critical temperature (CT_min_: this study CT_50/low_), upper critical temperature (CT_max_: this study CT_50/high_), thermal performance breadth (TPB) and mortality percentage estimates for the 49 F. columnare strains from Finland.

**Sup. Table S2. PCA summary**. The first five rows describe the matrix of loading.

